# Transkingdom Network Analysis (TkNA): a systems framework for inferring causal factors underlying host-microbiota and other multi-omic interactions

**DOI:** 10.1101/2023.02.22.529449

**Authors:** Nolan K. Newman, Matthew Macovsky, Richard R. Rodrigues, Amanda M. Bruce, Jacob W. Pederson, Sankalp S Patil, Jyothi Padiadpu, Amiran K. Dzutsev, Natalia Shulzhenko, Giorgio Trinchieri, Kevin Brown, Andrey Morgun

## Abstract

Technological advances have generated tremendous amounts of high-throughput omics data. Integrating data from multiple cohorts and diverse omics types from new and previously published studies can offer a holistic view of a biological system and aid in deciphering its critical players and key mechanisms. In this protocol, we describe how to use Transkingdom Network Analysis (TkNA), a unique causal-inference analytical framework that can perform meta-analysis of cohorts and detect master regulators among measured parameters that govern pathological or physiological responses of host-microbiota (or any multi-omic data) interactions in a particular condition or disease.

TkNA first reconstructs the network that represents a statistical model capturing the complex relationships between the different omics of the biological system. Here, it selects differential features and their per-group correlations by identifying robust and reproducible patterns of fold change direction and sign of correlation across several cohorts. Next, a causality-sensitive metric, statistical thresholds, and a set of topological criteria are used to select the final edges that form the transkingdom network. The second part of the analysis involves interrogating the network. Using the network’s local and global topology metrics, it detects nodes that are responsible for control of given subnetwork or control of communication between kingdoms and/or subnetworks.

The underlying basis of the TkNA approach involves fundamental principles including laws of causality, graph theory and information theory. Hence, TkNA can be used for causal inference via network analysis of any host and/or microbiota multi-omics data. This quick and easy-to-run protocol requires very basic familiarity with the Unix command-line environment.

## Introduction

Complex relationships between genetics and epigenetics are the basis of human health and disease. Advances in experimental and computational capabilities have brought a wide variety of high-throughput data to the study of biological systems. A large amount of technological effort is devoted to increasing throughput, reducing financial and personnel costs, and increasing experimental and computational efficiencies. As such, there is a clear interest in computational methods and software that can integrate different types of omics data and perform association analyses to identify important players and mechanisms (e.g., *mixOmics*). TkNA, however, is a unique causal-inference analytical framework that can perform meta-analysis and identify the regulatory relationships between entities. This approach, for example, was used in a study of antibiotic-resistant microbes^1^ to identify key host genes, microbes and molecular mechanisms in type 2 diabetes^2^. TkNA has also been used to characterize the role of the microbiome in cervical cancer, lymphoma and melanoma^3–6^. Please note that while the word “integration” refers to two very different analyses in other software, we have used it in the following way throughout the paper: meta-analysis is the *<u>integration of</u>* data from several independent *<u>datasets</u>*, whereas network reconstruction involves *<u>integration</u>* that establishes statistical dependencies between multiple types *<u>of omics data</u>*.

In this protocol, we describe in detail how to use TkNA to detect master regulators (causal factors) among microbes and microbial genes, host pathways and host genes, and other measured variables that govern pathological or physiological responses of host-microbiota interactions in a particular condition or disease. Note that although the TkNA approach was developed to investigate host-microbiota interactions, its underlying basis involves fundamental principles including laws of causality, graph theory and information theory. Hence, TkNA can be used for causal inference via network analysis of any multi-omics data collected whether those data were measured in the same samples or measured in different organs from the same biological replicates of animals (humans, mice, other animals) and/or environmental niches.

## Overview of the procedure

The TkNA pipeline consists of three major sections (**Figure 1**). **Section 1 reconstructs the network**. This involves importing the data, performing calculations/meta-analysis, and filtering the data as specified by the user. Specifically, differentially expressed/abundant variables (genes, microbes, metabolites, etc.) between classes (e.g., disease and control) are found, based on user-defined statistical criteria. Next, per-group correlations are performed within and between each type of omics/kingdom separately. To override the default comparison and correlation methods, appropriate parametric or non-parametric methods can be selected by the user. TkNA identifies robust and reproducible patterns of fold change direction (increased or decreased) and sign of correlation coefficient (positive or negative) across several cohorts (or experimental replicates). Following this, correlations are further filtered based on a causality-reflecting metric, correlation inequalities^7^, where within-class correlations with the opposite sign of coefficient than would be expected by the fold change direction of the variables in the edge are removed (i.e., if both variables of the edge are increased or decreased in the disease class compared to the control class, the expected per-group sign of correlation coefficient between those variables is positive and therefore negative correlations between them would be removed; for variables with opposite fold change direction in disease vs control the expected per-group sign of correlation coefficient is negative and therefore positive correlations between them would be removed). Finally, TkNA calculates a set of topological criteria, including network density, the deviation of observed positive:negative correlations from expected, and the proportion of unexpected correlations^7^ (PUC). The user employs these metrics to determine the quality of the reconstructed network by comparison to “typical” networks (networks we have reconstructed and published previously using a variety of omic data^5,8–13^). If the network is deemed suitable for downstream analysis, the user moves on to the network analysis step. Otherwise, the user changes their statistical thresholds and performs another reconstruction as described above. Of note, one of the output files of this step is a comma-separated value (CSV) file of the network, which the user can visualize using an external program such as Cytoscape^14^.

**Figure 1.**
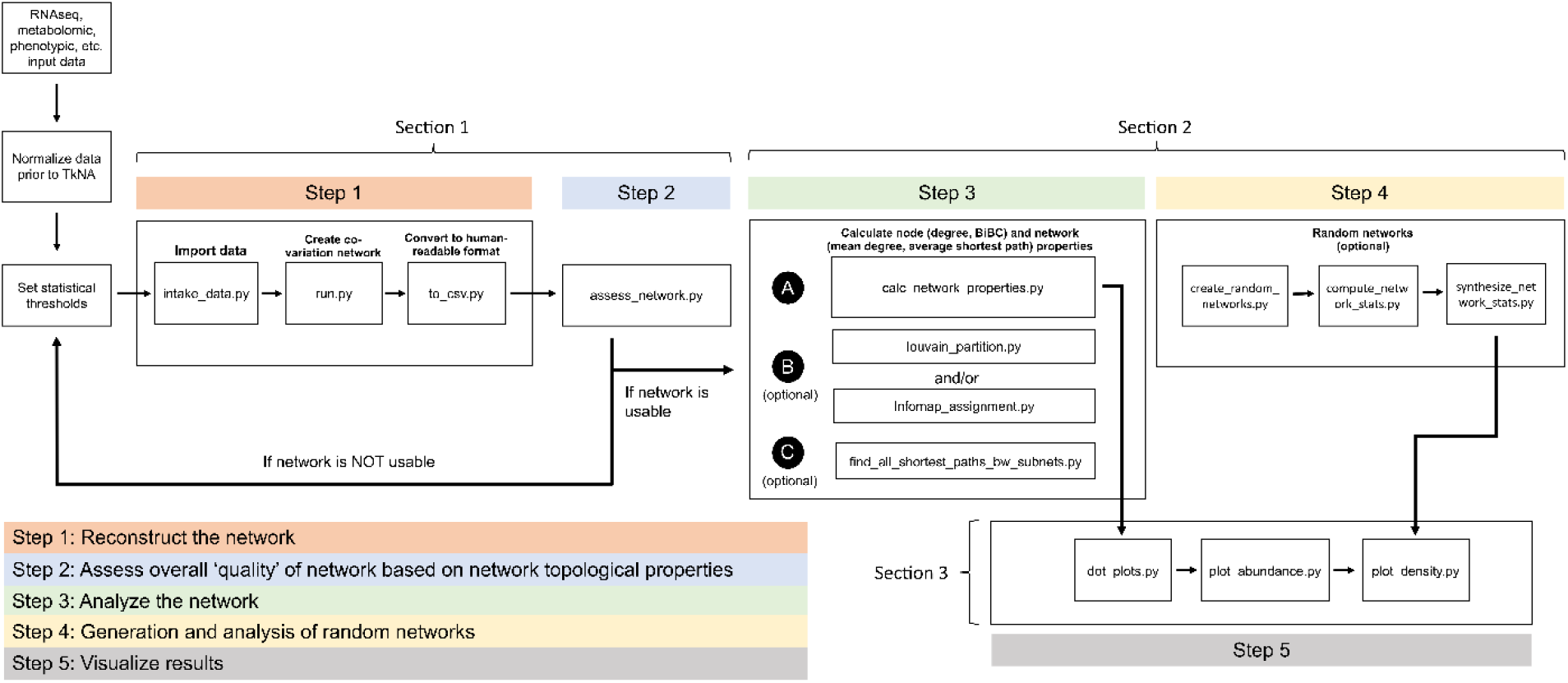
Flowchart of the TkNA pipeline.

**Section 2 is the interrogation/analysis of the reconstructed network**. Here, the user has the option to identify clusters of nodes in the network using the Louvain^15^ or Infomap^16^ algorithms. Using external recommended software (Table 3), they can then perform enrichment analysis of the clusters to identify biological pathways/functions to which the nodes in the cluster contribute. Regardless of whether cluster identification is performed, the next step in TkNA is to calculate node and network properties. While many different topological properties are calculated, key among these are node properties that indicate regulatory nodes in the network. This includes calculating bipartite betweenness centrality^17^ (BiBC), a global node topology metric, to find the nodes participating highest in the information flow between subnetworks or user-defined regions (e.g., microbe and host subnetworks) of the network. Degree is a key local node property that is calculated, which is a measure of how many other nodes to which a single node connects. High BiBC and high degree reflect “bottleneck” and “hub” nodes, respectively. Consequently, a node with a high degree and BiBC is considered to be a strong regulatory candidate in the network. Optionally, the user can also calculate shortest path lengths between each pair of nodes in two separate subnetworks. Since these calculations use the number of edges but not their strength (e.g., sign and magnitude of correlations), two subnetworks with a smaller average shortest path length are predicted to interact more (but not necessarily more strongly) than subnetworks that are farther away from one another. To wrap up section 2 and evaluate the probability of a given node to show nonrandom values of degree and BiBC, TkNA reconstructs many random networks (10,000 by default) with the same number of nodes and edges as the reconstructed network and compares the top degree/BiBC nodes of those random networks to the reconstructed network.

**Table 1.**
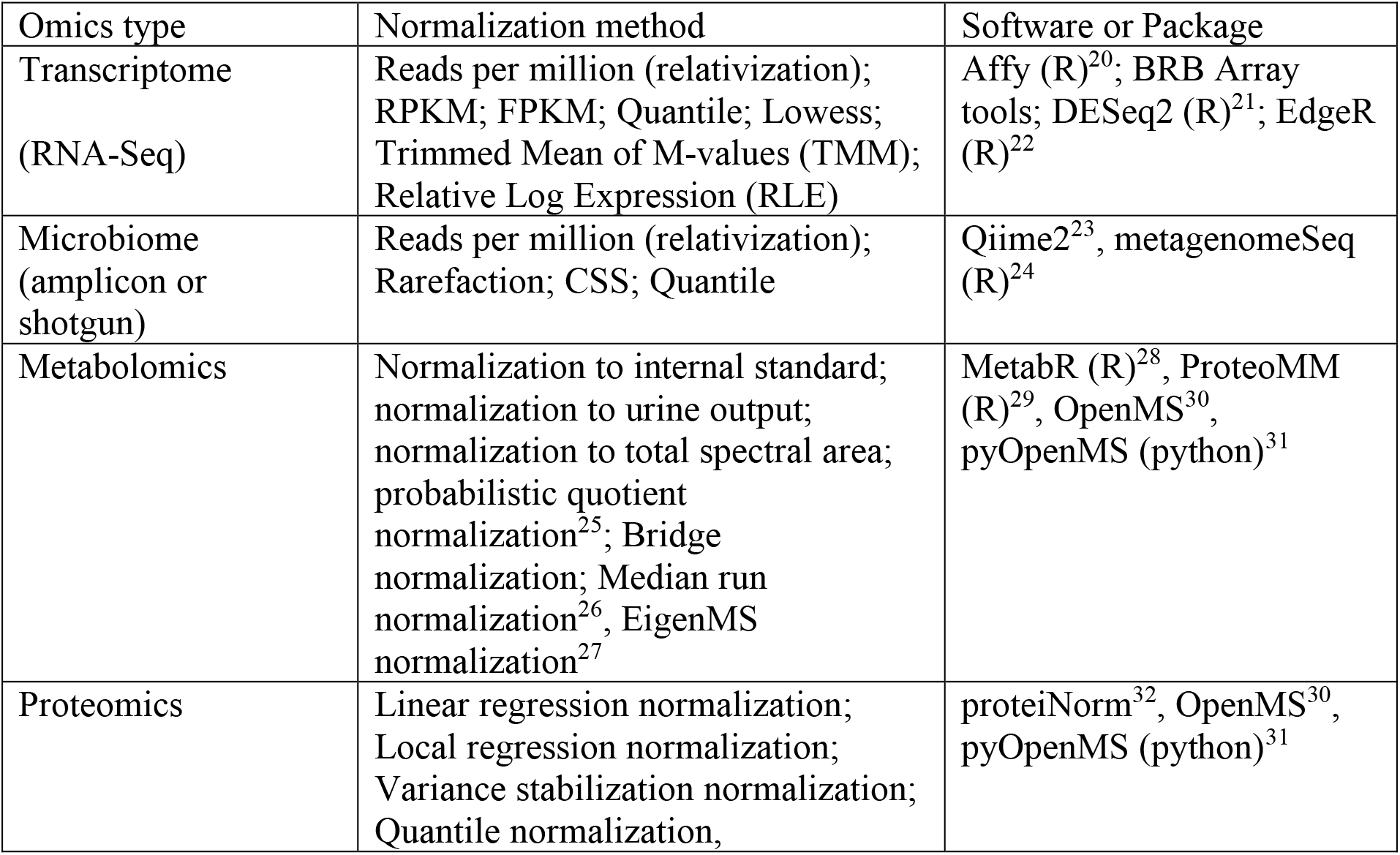
Common normalization methods for various omics types.

**Table 2.** Suggestions for alterations in thresholds based on the resulting network assessment properties.

| Network property | Potential range | If value is too high... | If value is too low... |
| --- | --- | --- | --- |
| Departure of ratio positive:negative edges | See figure 4a | Try alternative normalization | Try alternative normalization |
| Density | Dependent on -omic type, see figure 4b | Reduce statistical thresholds (p-values or FDR) for edges | Relax statistical thresholds (p-values or FDR) for edges |

**Table 3.**
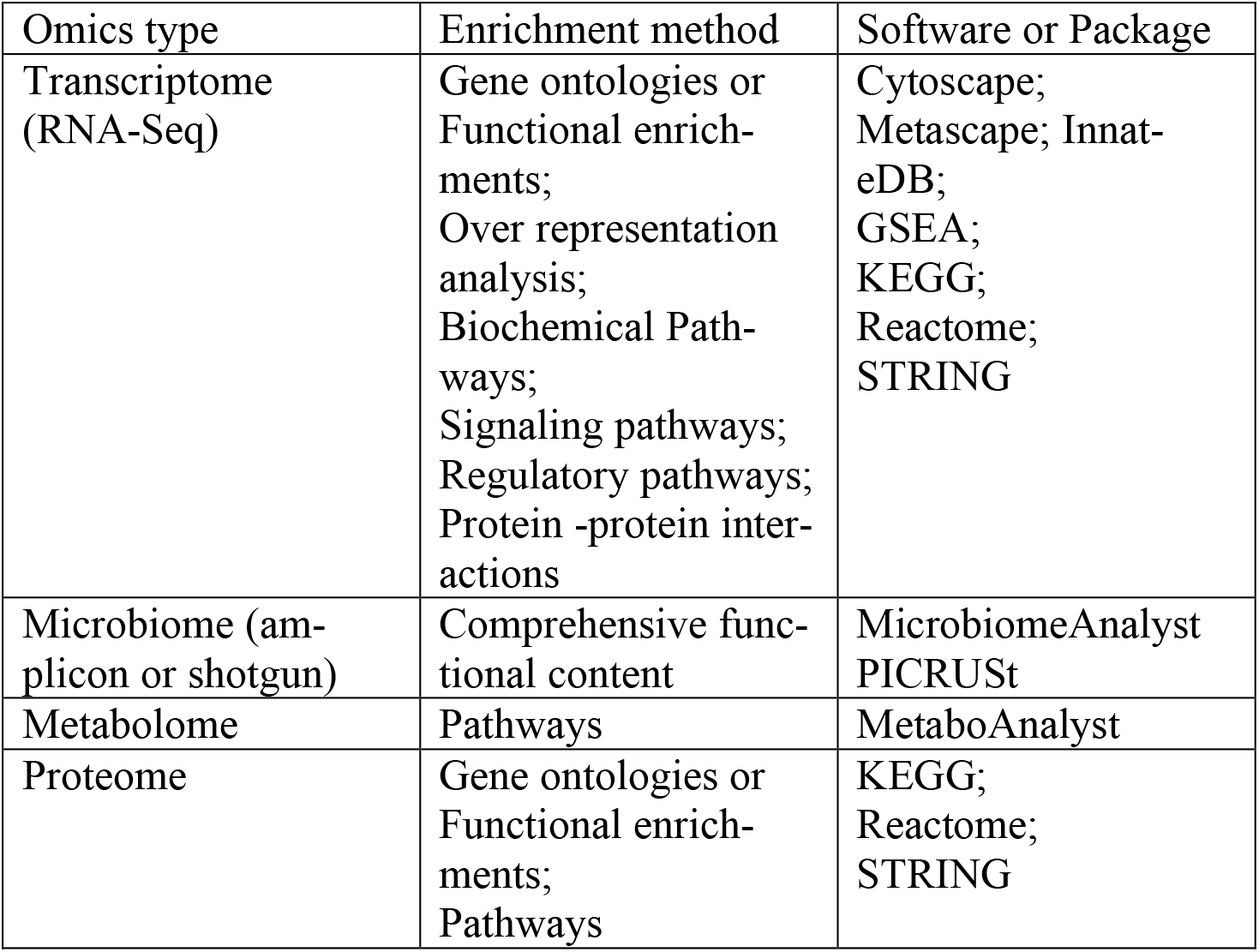
Examples of common functional enrichment methods for different omics data.

**Section 3 of TkNA creates publication-ready figures from the above analysis of the user’s reconstructed network**. Multiple high-quality figures are created in this step, including dot plots of degree distribution (**Figure 7a**) and dot plots of nodes and their calculated properties (**Figure 7b**). The abundance (e.g., microbiome) or expression levels (e.g, transcriptome), etc. of the top regulatory candidates are also automatically generated (**Figure 7c**), as well as a 2D density plots (**Figure 7d**) that have the observed values of the top regulatory nodes from the reconstructed network overlayed on top of them. Additionally, CSV files are created with all the necessary information if the user wishes to use an alternative visualization or plotting software.

**Figure 2.**
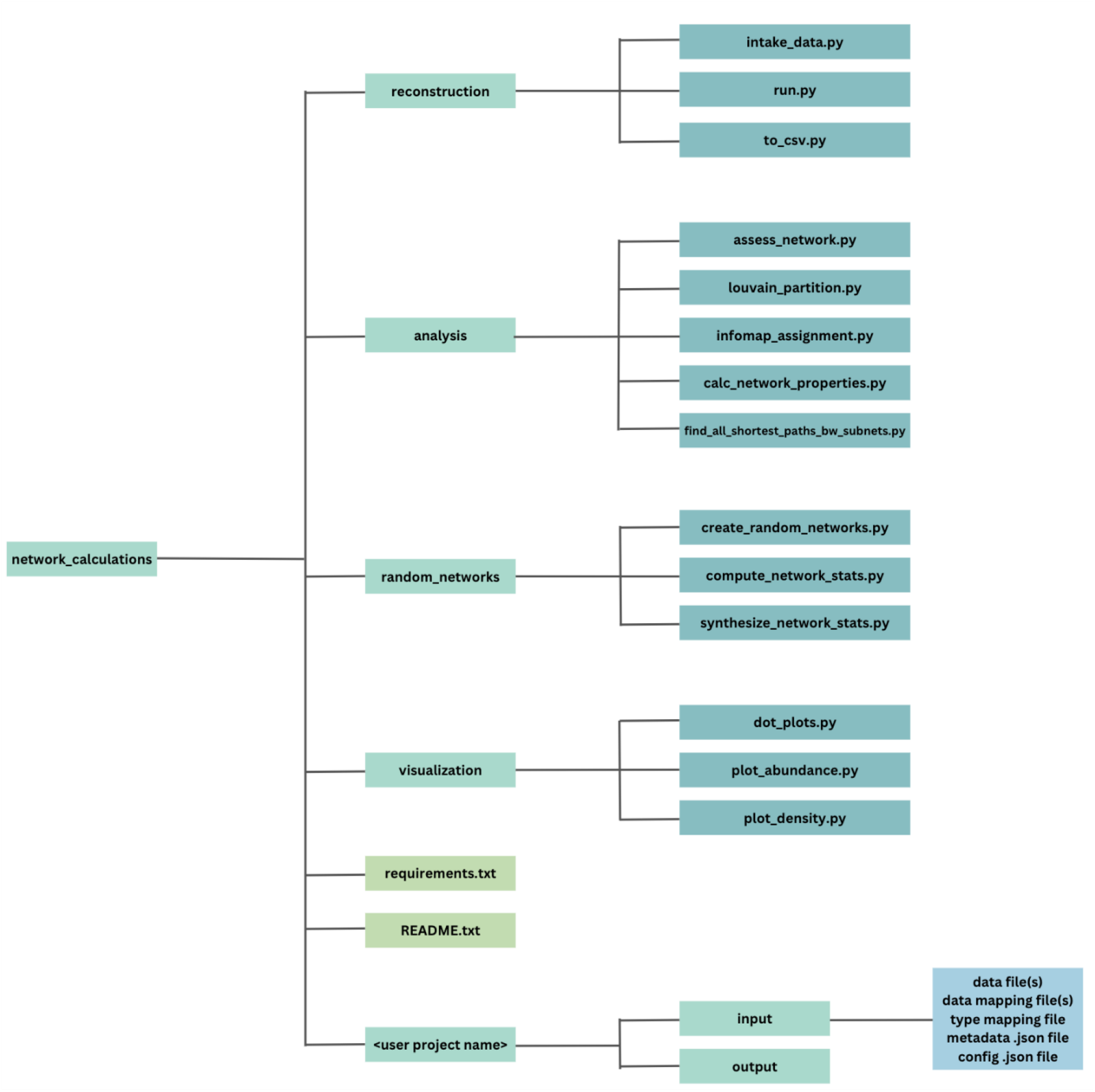
Structure of the TkNA repository, including all folders and the main scripts required for the user to reference in their commands. We recommend the user creates a directory for their project like the one at the bottom of the figure, where all the input files are stored and where they will direct the output files to write to in their commands.

**Figure 3.**
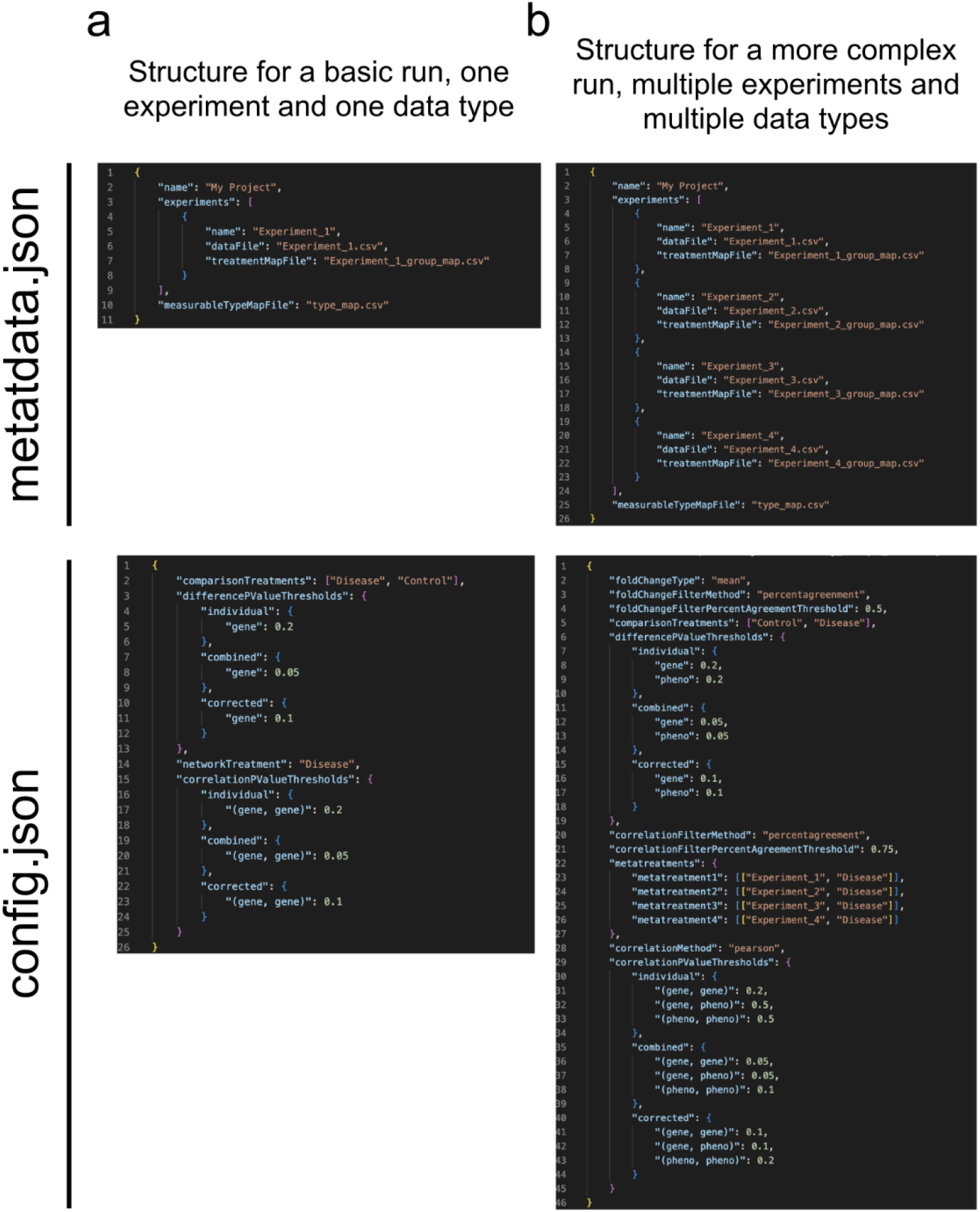
Example metadata and configuration (config) .json files. **a)**The structure required to run the network reconstruction code for a single dataset. **b)**The structure required for more complex data with multiple experiments and/or data types.

**Figure 4.**
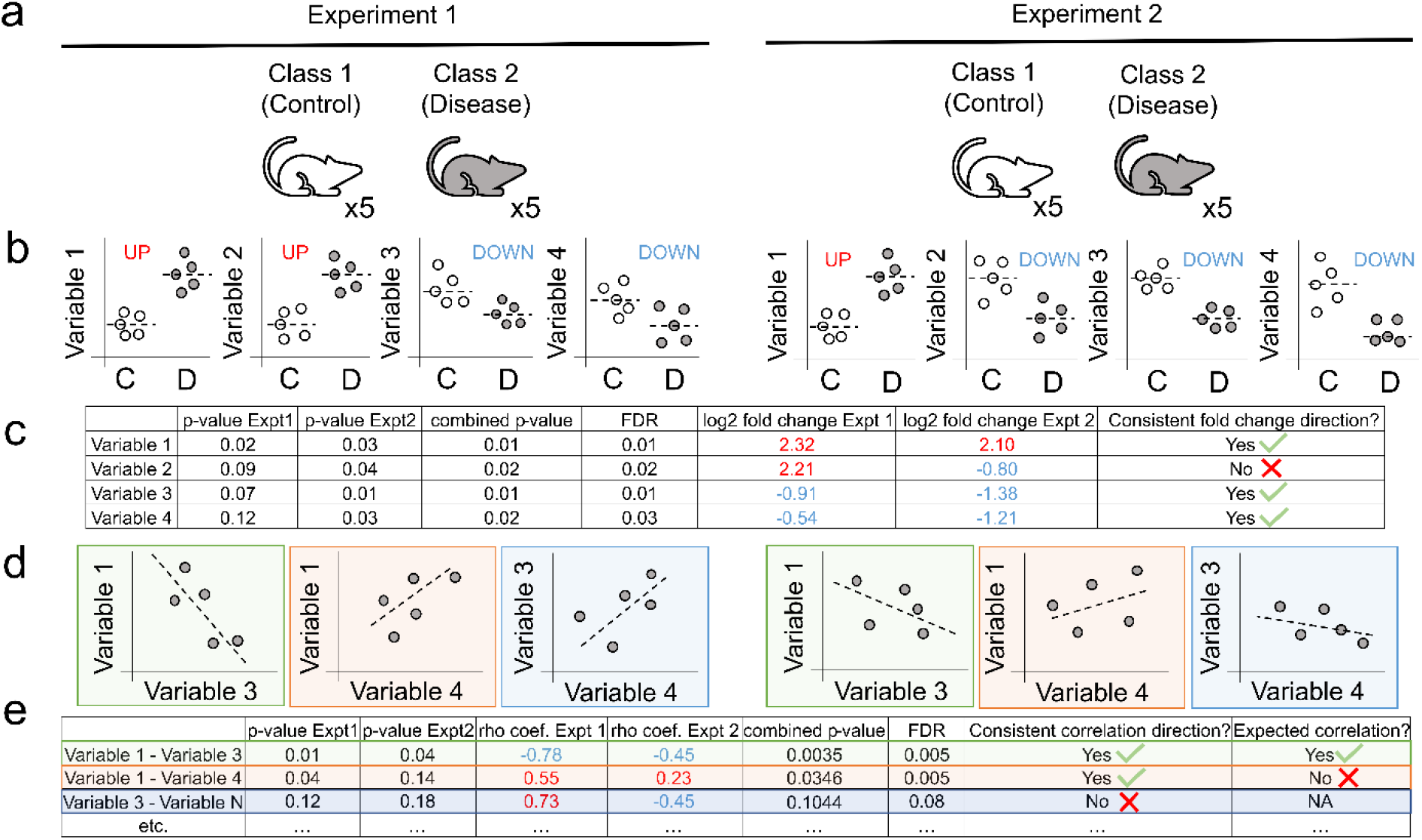
Schematic of how a basic network is reconstructed from two independent datasets. a) An experimental design that works well for the TkNA pipeline, allowing for meta-analysis with robustness of findings across experiments. Note that a minimum of five samples per class per experiment is recommended. b) Comparisons of each variable between the two classes (defined by user) are performed. A p-value (either from a Student’s t-test or Mann-Whitney test) and fold change are calculated for each variable in each experiment. Meta-analysis is then conducted by calculating the Fisher’s combined meta-analysis p-value and FDR. c) The variables are filtered to remove those that did not pass the user-defined p-value, combined p-value, or FDR. Additionally, only variables that are the same direction of fold change across the two experiments are retained. d) Following the filtering in c, correlations between the remaining variables are performed per-group. A p-value (either Spearman or Pearson) and rho coefficient are calculated for each pair of remaining variables. Meta-analysis is then conducted by calculating the Fisher’s combined meta-analysis p-value and FDR. e) Correlations are filtered and those that did not pass the user-defined p-value, combined p-value, or FDR are removed. Additionally, correlations that are not consistent in direction (positive or negative) are removed. Unexpected correlations (a positive correlation between two nodes of opposite fold change direction or a negative correlation between two nodes of the same fold change direction) are also removed. The reconstructed network is comprised of the remaining nodes and edges.

**Figure 5.**
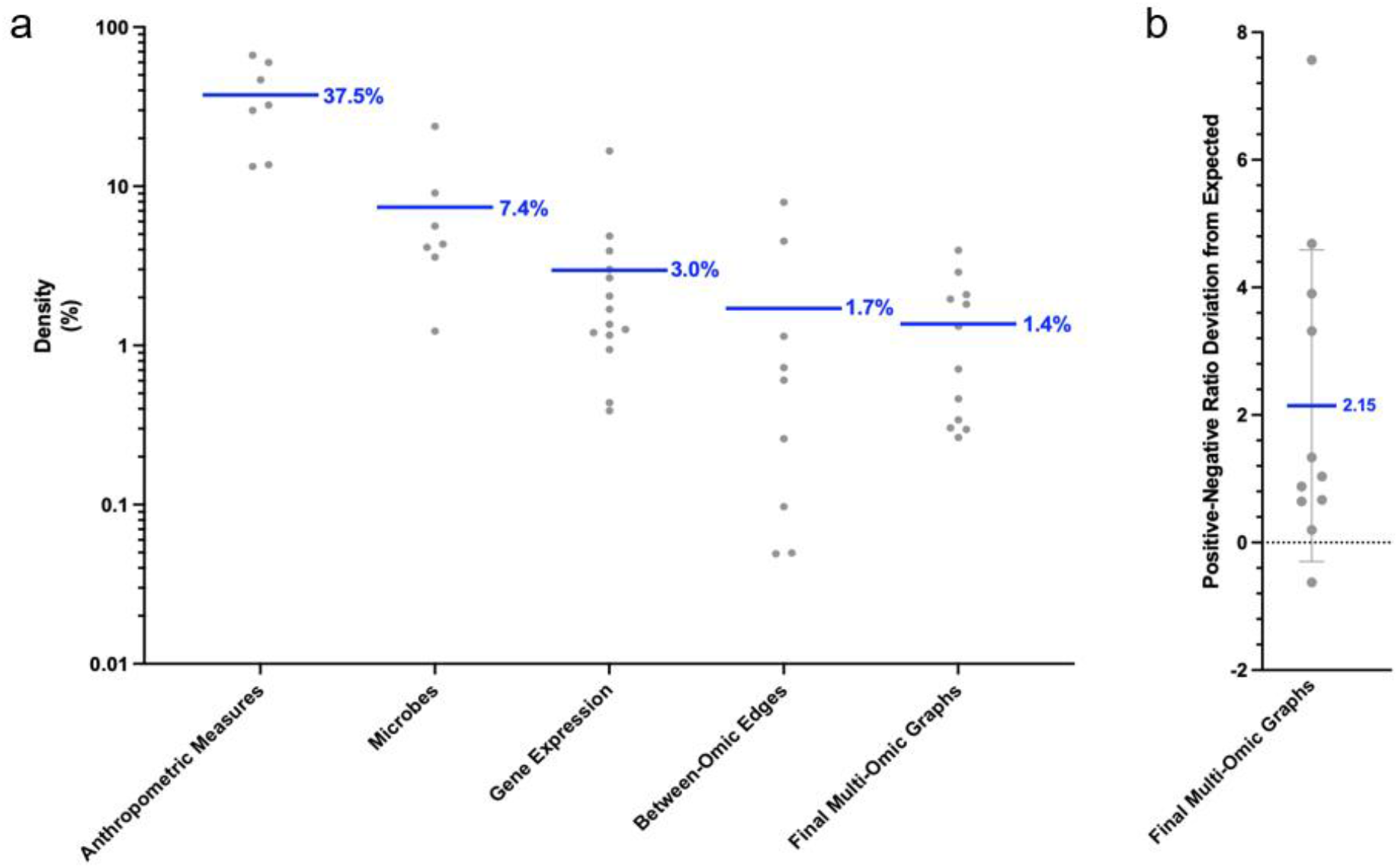
**a)** Density of within-omic and between-omic edges in published networks. Each point represents a previously published network. Density is calculated as the number of observed edges over the number of possible edges between all nodes of that type. The blue bar indicates the average of each –omic type. **b)** The deviation from the expected positive/negative edge ratio of published networks. Each point represents a previously published network. The blue bar indicates the average.

**Figure 6.**
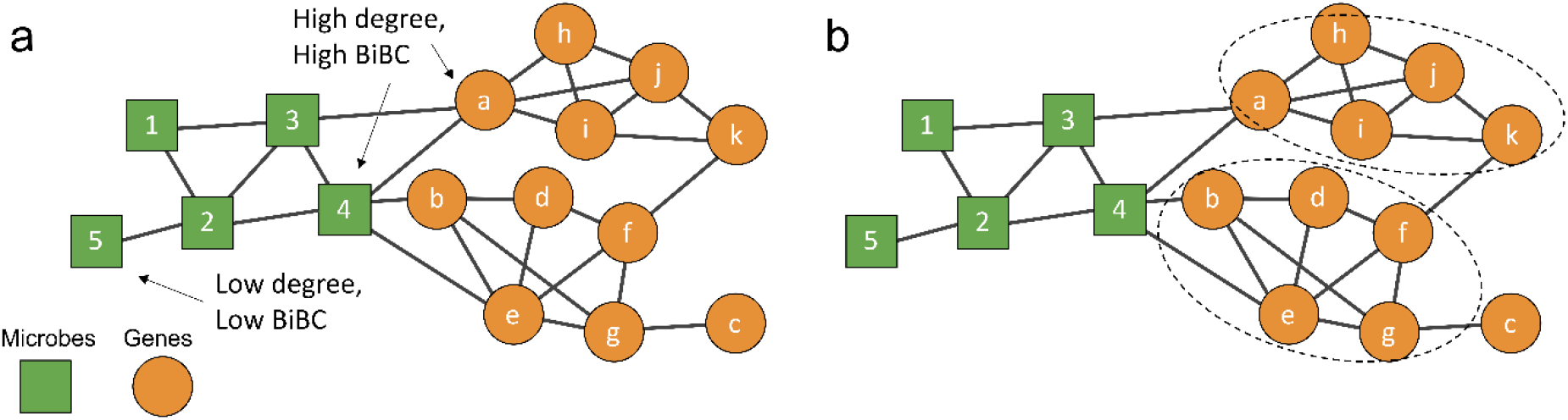
a) Examples of nodes in a network that are not regulatory (low degree and low BiBC) and nodes that are regulatory (high degree and high BiBC). Squares are microbes and circles are genes. b) Example of clustering in the same network using the Louvain or Infomap algorithms.

**Figure 7:**
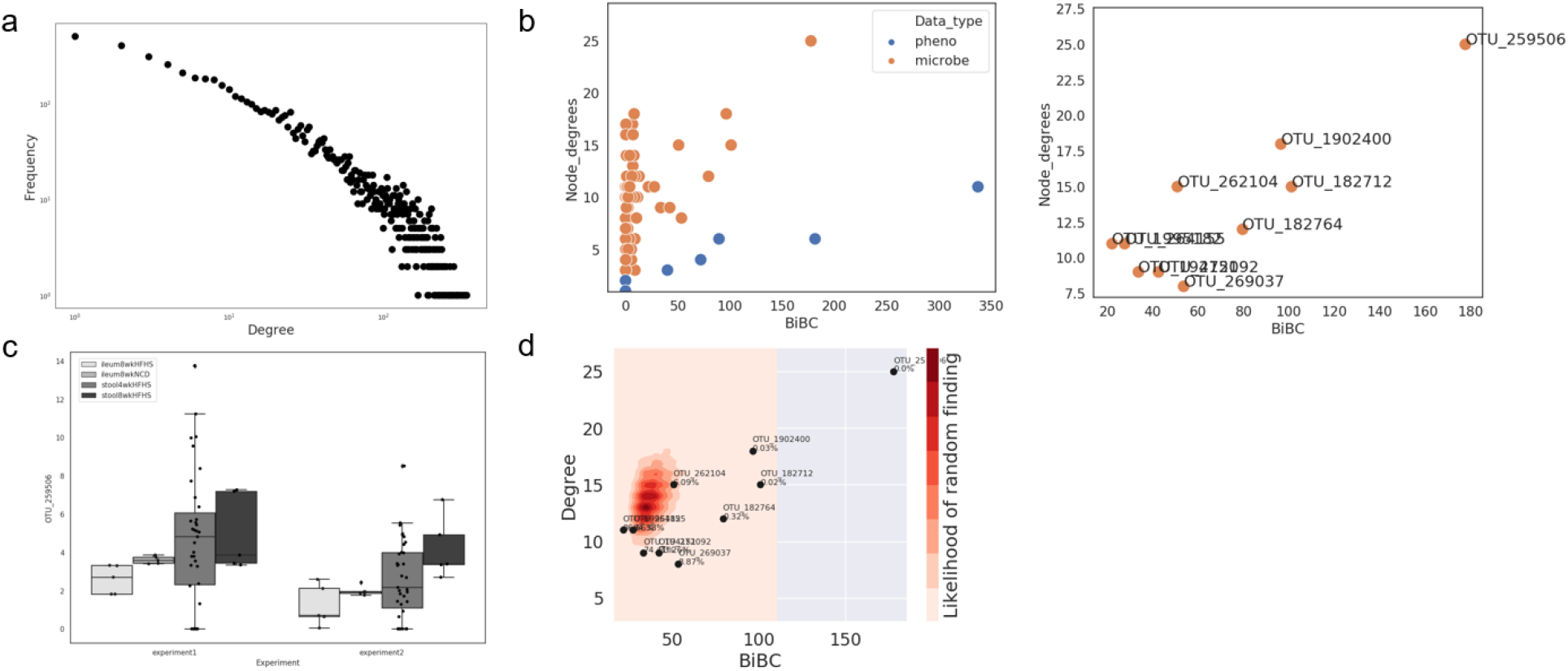
**a)** Degree distribution plot **b) Left:** Example node property visualization, where each point represents a node in the reconstructed network. **Right:**Same graph, zoomed in on the top 5 microbe BiBC nodes **c)** Example abundance/expression plot of the top BiBC node found in b. Legend is the two classes that were in the dataset. In this case, a class of samples named ‘high’ was compared to a class named ‘low’ **d)** 2D density plot of the 10,000 random networks with the top nodes from b overlaid.

## Procedure

### Pre-requirements for using TkNA

Currently, TkNA is only supported on Unix (e.g., Mac, Linux) based devices. From a Unix device, a user will need to have access to git on their terminal and have an SSH key for their GitHub account (see https://docs.github.com/en/authentication/connecting-to-github-with-ssh/adding-a-new-ssh-key-to-your-github-account for more information). If the user does not already have git downloaded, he or she can do so using the commands found on github’s documentation at https://github.com/git-guides/install-git.

## Software Setup

### 1. Obtain TkNA

Download the TkNA code from https://github.com/CAnBioNet/TkNA.git and enter its directory.

~~~
  git clone git@github.com:CAnBioNet/TkNA.git
  cd TkNA
~~~

### 2. Install Miniconda

We highly recommend that users manage their installation using conda as their package management software. If it is not already installed on the system, you can install Miniconda (which contains both python and conda) from https://docs.conda.io/en/latest/miniconda.html. Since TkNA was written and tested in Python version 3.8.10, we recommend the user installs the Python 3.8 Miniconda version. Computationally proficient users can run TkNA with other python versions, however, installing and troubleshooting with python versions other than v3.8.10 is outside the scope of this protocol, and it is upon the user to install the required libraries.

### 3. Set up the TkNA environment

Create a new Python 3.8.10 conda environment. Install the python packages specified in the `requirements.txt` file included with the source code.

~~~
  conda create -n TkNA python=3.8.10
  pip install -r requirements.txt
~~~

## Data Preprocessing

### 4. Normalize data

It is known that non-normalized data leads to bias in the structure of correlation networks^18,19^. As such, prior to running TkNA, the user needs to perform any appropriate normalization(s) for their dataset. Normalization is not performed in TkNA, so data can be normalized using but not limited to the methods listed in **Table 1**.

### 5. Format data

Following normalization, we recommend the data be organized into a directory and arranged as shown in **Figure 2**. After cloning the repo, the user will see all folders on their terminal, then will need to manually create a project folder for each project, with a folder to hold the input and output files inside.

Examples of each input files can be found in the repo at https://github.com/CAnBioNet/TkNA/tree/main/example_datasets_and_commands/microbiome_and_phenotype/input (referred to below as microbiome_and_phenotype/input/ for simplicity). There are five main file types required to run the TkNA pipeline:

a. **Processed (normalized and log2 transformed) data tables** Each data table associated with an experiment present in the data must be log2 transformed prior to the pipeline. Pseudocounts can be added to all values to avoid negative numbers after log transformation. The data table must be formatted as in comma-separated value (CSV) format with one column per experimental unit (i.e., sample) and one row per variable (e.g., a gene, microbe, or phenotype). Experiment1.csv and Experiment2.csv in the microbiome_and_phenotype/input/ folder on github are examples of these files. The variable names must not contain any characters other than letters, numbers, underscores, and spaces. If the user’s data contains any other characters such as commas, we recommend giving them a new unique alphanumeric ID and using a separate reference file to keep track of their new and old names.
b. **Sample mapping file** Each experimental unit present in the data must be associated with an experimental class (e.g. Disease, Control), identified by a text string. These associations are specified as lists of comma-separated values, with the left column containing the name of the experimental unit and the right column containing the class. All names must match exactly across data files. Currently, TkNA is designed to only work with two classes at a time. Note: In the pipeline, the sample mapping file is called “treatmentMapFile” in the metadata .json file (**Figure 3a**) and the sample groups to compare are the ones used in “comparisonTreatments” in the config .json (**Figure 3b**) file. Experiment1_group_map.csv in the microbiome_and_phenotype/input/ folder on github is an example of this file.
c. **Omics type mapping file** Each variable (e.g., microbe, gene, metabolite, etc.) present is associated with a type of omic data, identified by a text string. These associations are specified as lists of comma-separated values, with the left column containing the variable and the right column containing its type. All names must match exactly across data files. type_map.csv in the microbiome_and_phenotype/input/ folder on github is an example of this file.
d. **config file** The dataset must contain a file named `config.json` in the JavaScript Object Notation (JSON) format, as shown in **Figure 3**. This file defines the statistical and meta-analysis criteria to apply when reconstructing the network. More in-depth information and examples for this file can be found in **Step 7 - Reconstruct network**. config.json in the microbiome_and_phenotype/input/ folder on github is an example of this file.
e. **Metadata file** The dataset must also contain a file named `metadata.json` in JSON format, as shown in **Figure 3**. This file describes each experiment present in the data and indicates the names and locations of the necessary files. As such, the names in this file <u>must match exactly</u> the names of the files being used in the analysis. metadata.json in the microbiome_and_phenotype/input/ folder on github is an example of this file. At the top-most level, the metadata consists of 3 sections:

i. **Project name** A name to which the project will be referred by.
ii. **Experiment/Cohort/Dataset(s) details** A list of experiments/cohorts/datasets (which we will refer to as “experiments”) whose data is present in the project. Each experiment contains a “name”, the relative path to the CSV file containing its data table (“datafile”), and the sample mapping file name (“treatmentMapFile”) (**Figure 3a**).
iii. **Type map** The omics type mapping file “measurableTypeMapFile” for all of the variables present in the data. In **Figure 3a**, the users have a single experiment in their project (as seen in the metadata file) with two classes they would like to compare: Disease and Control (as seen in the config file). They have then specified which comparison statistical thresholds to apply (“differencePValueThresholds”) for individual p-value, Fisher’s combined (meta-analysis) p-value, and the Benjamini-Hochberg FDR (called “corrected” in the config file). Additionally, they have specified they want to perform the correlations only in the Disease class (“networkTreatment”) and specified the edge statistical criteria (“correlationPValueThresholds”) towards the bottom of the file. The user for **Figure 3b** has a more complicated project, with four total datasets, as well as an additional “pheno” variable type in their input data (genes and phenotypes), so they have added the additional “pheno” variable to the file. They have set the same comparison thresholds for both variable types, although matching thresholds are not required. Down below, they have specified they want to find only edges that are consistent in the Disease class of three of the four experiments (as can be seen by the 0.75 for “correlationFilterPercentAgreementThreshold” and the “metatreatments” sections). Then they specified the edge thresholds for each pair of data types (gene-gene, gene-pheno, and pheno-pheno). Please note that filtering edges for the same direction across only a fraction of datasets/groups is not typically a good strategy, especially when using a small number of datasets/groups (less than five) but was done in this example for simplicity. We recommend using “correlationFilterPercentAgreementThreshold” option as a last resort and only by expert users. In other words, the default of “correlationFilterMethod” keeps edges that are consistent across all datasets/groups (i.e., the same sign of correlations). This is critical to avoid situations where an edge’s direction is decided by most of datasets/groups with non-significant correlation coefficients instead of the fewer datasets/groups with strong significant correlation coefficients but with opposite sign to those in the majority of the datasets/groups. Also note that the structure of .json files is highly specific and the addition of extra characters in the files may cause issues.

## Data Import

### 6. Import data

Once all the required files have been uploaded, the next step is to import the data into a consolidated format for downstream processing, using the script `reconstruction/intake_data.py`.

**Usage**

~~~
    python ./reconstruction/intake_data.py --data-dir <data
    directory> --out-file <output file>
~~~
**Example command**

~~~
    python reconstruction/intake_data.py --data-dir
    ./project_folder/input/ --out-file
    ./project_folder/output/all_data_and_metadata.cdf
~~~
**Inputs**

`--data-dir`: Path to the directory containing all experimental file(s), metadata file(s), and config file(s)
`--out-file`: path to file (with .cdf extension) that will be created
**Outputs**

- A single .cdf file containing most information required for the next step

## Network Reconstruction

### 7. Reconstruct network

Network reconstruction is then performed with the script `reconstruction/run.py`, which references the JSON configuration file (**Figure 3**). An overview of the network reconstruction process can be found in **Figure 4**.

**Usage**

~~~
    python ./reconstruction/run.py --data-source <file_name> --
    config-file <config file> --out-file <zip directory>
~~~
**Example command**

~~~
    python ./reconstruction/run.py --data-source
    ./project_folder/output/all_data_and_metadata.cdf --config-file
    ./project_folder/input/config.json --out-file
    ./project_folder/output/network_output.zip
~~~
**Inputs**

`--data-source`: Path to the .cdf file created using `intake_data.py`
`--config-file`: Path to the config file used for `intake_data.py`
`--out-file`: path to zipped directory that will be created
**Outputs**

- A single zipped directory containing the analysis performed

Reconstruction takes place in the following steps. For each step, the available configuration options are described. For more specific examples of the config file, please refer to **Figure 3**.

i. **Establish differentially expressed/abundant variables (genes, microbes, metabolites, etc.) between classes (e.g., disease and control).** Setting the appropriate statistical thresholds to apply while reconstructing the network is the most critical step in determining the size and quality of the resulting network. Thus, it is crucial that the user sets statistical thresholds that are best suited for the type of data they are using. For finding differentially expressed/abundant variables between classes, we recommend that the user initially uses the default thresholds supplied in the code: p-value < 0.2, Fisher’s combined (meta-analysis) p-value < 0.05, and Benjamini-Hochberg FDR <0.1. If the reconstructed network is not of sufficient quality (see **step 9 “Assessing network quality”**) or size (an experienced user may consider the network too small) then the user can select different thresholds to apply. At this stage of the analysis, it is not recommended to relax thresholds more than the following: individual p-value, 0.5; Fisher’s combined p value, 0.1; and FDR, 0.2. Note that relaxed thresholds can be used at this stage because a considerable proportion of variables will be further eliminated in the next step of the analysis when edges are established. **Required config parameter:** `comparisonTreatments`

A list containing the names of exactly two groups. Fold change is computed with respect to the first of the two, meaning “control” would be the denominator in the below example.
Example: `“comparisonTreatments”: [“control”, “disease”]` **Optional config parameter:** `differenceMethod`

The method to use for calculating correlations, either Mann-Whitney or Student’s t-test (for data with equal variance).
Choices: `mannwhitney, independentttest`
Default: `mannwhitney`
Example: `“differenceMethod”: “mannwhitney”` **Optional config parameter**: `differencePValueThresholds` A mapping between each kind of statistic and its threshold. Can contain:

individual: Minimum p-value across all experiments

Default: 0.2
combined: p-value combined across experiments (e.g., via Fisher’s method)

Default: 0.05
corrected: p-value after correction (e.g., via FDR)

Default: 0.1 NOTE: We strongly recommend using more than one dataset for the TkNA analysis. However, if the user only has one dataset, the combined p-value is the same as the individual p-value and the same threshold value should be used for both. Example: `”differencePValueThresholds”: {“individual”: 0.1, “combined”: 0.02, “corrected” 0.15}` Optionally, for each kind of statistic, thresholds can be specified per type of variable. Example: `“differencePValueThresholds”: {“corrected”: {“gene”: 0.1, “phenotype”: 0.15}}`
**Identify robust and reproducible patterns for fold change direction across several experimental replicates (if user has multiple independent experiments)** **Optional config parameter :** `foldChangeType`

Calculate fold change using the median or the mean.
Choices: `median, mean`
Default: `median`
Example: `“foldChangeType”: “mean”` **Optional config parameter :** `foldChangeFilterMethod`

Choose how strictly the fold changes of variables must be consistent across experiments.
Choices: `allsamesign, percentagreement`
Default: `allsamesign`
Example: `“foldChangeFilterMethod”: “percentagreement”`
If using percent agreement, the following option may also be specified: `foldChangeFilterPercentAgreementThreshold`

A fraction of experiments which must agree in fold change direction for a variable to pass the filter. While full concordance would mean 1.0, when relaxed thresholding is needed we recommend using values of 0.75 or higher.
Example: “foldChangeFilterPercentAgreement”: 0.8
**Determine significant correlations for a treatment within and between each type of variable (e.g., gene-gene, gene-microbe, microbe-microbe, etc.) separately.** To calculate correlations between the previously identified differentially expressed/abundant variables between classes, we recommend that the user initially uses the default thresholds supplied in the code: p-value < 0.2, Fisher’s combined (meta-analysis) p-value < 0.05, and Benjamini-Hochberg FDR < 0.1. At this stage of analysis, it is not recommended to relax thresholds more than the following: individual p-value, 0.5; Fisher’s combined p value, 0.05; and FDR, 0.15. The groups of samples from which a network is reconstructed can be specified in two ways. The simplest way is to specify the name of a single class (e.g., disease) for networkTreatment. In this case, correlations will be calculated within this group for each experiment, then meta-analysis will be performed between the experiments. If one wants to include samples from different classes in the meta-analysis, the parameter `metatreatments` can be specified instead. See below for more details on the use of `metatreatments`, and see section “**Experimental Design**”, as well as **Figure 8**, for details on when to apply them and in what manner. If you are reconstructing the network from a single class (either in one experiment or across multiple experiments):

**config parameter :** `networkTreatment`

The name of a class to produce a network from. Either this or metatreatments must be specified.
Example: `“networkTreatment”: “disease”` Otherwise, if you are reconstructing the network from multiple different groups (e.g., two different disease groups from one experiment and another disease groups from a second experiment):

**config parameter:** `metatreatments`

A description of “metatreatments” to use for creating the network. Either this or `networkTreatment` must be specified.
Each metatreatment consists of (experiment, class) pairs. Correlations are calculated for each metatreatment using all of the data specified for it. Meta-analysis will then be performed across all of the specified metatreatments.
Example:

~~~
  “metatreatments”: {
  “metatreatment1”: [[“experiment_1”, “disease_1”]],
  “metatreatment2”: [[“experiment_2”, “disease_1”]],
  “metatreatment3”: [[“experiment_2”, “disease_2”]]
  }
~~~ The above example corresponds to meta-analysis option #3 in **Figure 8**. Note that using “correlationFilterPercentAgreementThreshold” along with this option can have unintended consequences as discussed previously (e.g., two non-significant positive correlations from disease 1 could win over one significant negative correlation from disease 2, marking the edge to be positive instead of being discarded as would have happened with the default threshold of “correlationFilterMethod”: “allsamesign”). **Optional config parameter:** `correlationMethod`

The method to use for calculating correlations.
Choices: `spearman, pearson`
Default: `spearman` **Optional config parameter:** `correlationPValueThresholds` A mapping between each kind of statistic and its threshold. Can contain:

individual: Maximum raw p-value across all experiments

Default: 0.2
combined: p-value combined across experiments (e.g., via Fisher’s method)

Default: 0.05
corrected: p-value after correction (FDR via Benjamini-Hochberg)

Default: 0.1
Example: `“correlationPValueThresholds”: {“individual”: 0.1, “combined”: 0.02, “corrected” 0.15}`
Optionally, for each kind of statistic, thresholds can be specified per combination of types of variables.
Example:

~~~
  “correlationPValueThresholds”: {“corrected”: {
  “(gene, gene)”: 0.1,
  “(gene, phenotype)”: 0.15,
  “(phenotype, phenotype)”: 0.2
  }}
~~~
**Identify robust and reproducible patterns for correlation direction across several cohorts (or experimental replicates).**

**Optional config parameter:** `correlationFilterMethod`

Choose how strictly the correlations of variables must be consistent across experiments.
Choices: `allsamesign, percentagreement`
Default: `allsamesign`
Example: `“correlationFilterMethod”: “percentagreement”`
If using percentagreement, the following option may also be specified: `correlationFilterPercentAgreementThreshold`

A fraction (greater than 0.75 and less than 1) of experiments which must agree in sign of correlation coefficient for a variable to pass the filter.
Example: `“correlationFilterPercentAgreementThreshold”: 0.8`
ii. **Eliminate edges that are not related to causes underlying the transition between classes (e.g., between disease and control)** This step is performed automatically. The edges removed are defined as “unexpected correlations”^7^. The edges labeled as unexpected can be found in the correlations_bw_signif_measurables.csv file.

**Figure 8.**
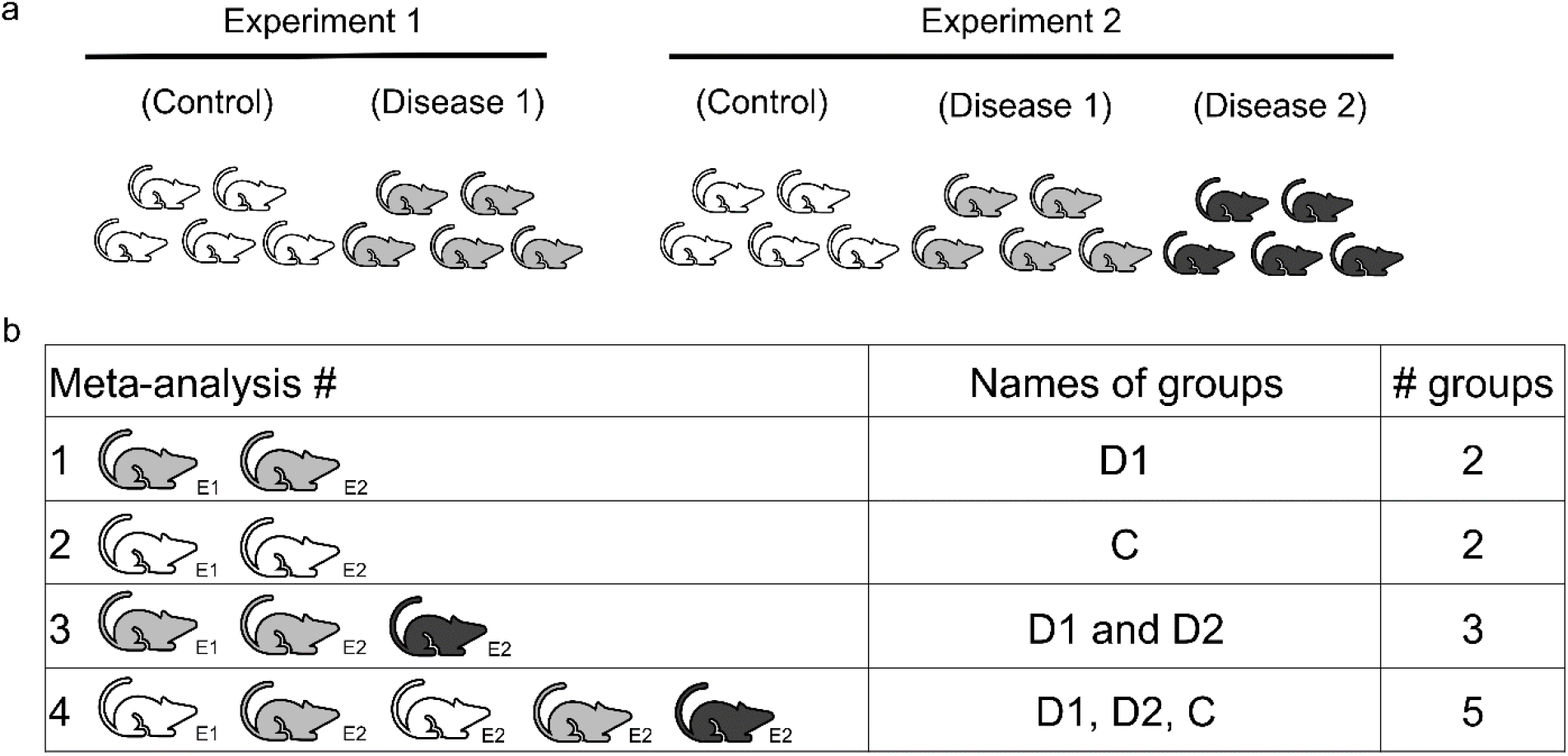
Example configurations of experimental classes for meta-analysis of edges. **a)**An example study, in which the user has two experiments. In experiment 1, there are two classes: control and disease. In experiment 2 there are three classes: the same control class as experiment 1, the same disease class as experiment 1, and a second disease class, not present in experiment 1. **b)**Different options for meta-analysis. Option 1: The ideal (recommended) option for meta-analysis, in which the same disease groups from both experiments are used. Option 2: Same as option 1 except uses the control groups. Option 3: Use all three disease groups to find the underlying edges that are present in the disease, regardless of potential differences between D1 and D2. Option 4: Use all five groups. This option presents the highest statistical power and generalizability, finding the edges present across all conditions (classes).

### 8. Write computed data to tables

Write the output to tables in CSV format using the script `reconstruction/to_csv.py.`

**Usage**

~~~
  python ./reconstruction/to_csv.py --data-file <zip file> --
  config-file <config file> --out-dir <output directory>
~~~
**Example command**

~~~
  python ./reconstruction/to_csv.py --data-file
  ./project_folder/output/network_output.zip --config-file
  ./project_folder/input/config.json --out-dir
  ./project_folder/output/network_output
~~~
**Inputs**

`--data-file`: .zip file created with `run.py`
`--config-file`: Path to the config file used for `intake_data. py`
`--out-dir`: Path to the directory to output results to
**Outputs**

- `all_comparisons.csv`: contains all comparisons performed in the analysis, is not filtered for any significant variables.
- `correlations_bw_signif_measurables.csv`: all correlations between variables that passed the comparison thresholds. Correlations in this file are not filtered for statistical or causality criteria, but it contains all p-values, whether each edge is unexpected, and whether each edge makes it into the final network after applying statistical and causality criteria.
- `network_output_comp.csv`: whole network with nodes/edges under the user-defined statistical thresholds, and edges consistent in direction retained (unless otherwise specified in the configuration file). Unexpected edges are also removed from this file.
- `node_comparisons.csv`: comparisons that were performed and found to be statistically significant (less than the user-defined statistical thresholds).
- `config_values.txt`: All the user-specified options for making the network.

## Network quality assessment

Once the network has been reconstructed, the next step is to determine whether the network is suitable for downstream analysis. If the network is not suitable for downstream analysis, the user will need to modify their statistical thresholds to reconstruct a better-suited network.

While as yet there is no completely automated method of network optimization, we recommend the user evaluates three network properties that are calculated with TkNA: 1) the proportion of unexpected correlations (PUC); 2) density (the number of edges in the network over the number of edges in a full graph of the same size); and 3) the deviation of the ratio of positive/negative edges from the expected value. In **Table 2** we provide the ranges of each of these criteria from previously published networks that can guide the user’s decision process. Note that microbiome data usually presents higher PUC values than diverse host omics data (e.g., transcriptomes, metabolomes, etc.). The user can also compare their values with those of published networks using **Figure 5**.

We encourage new users to use the **<u>default statistical thresholds</u>** for reconstructing their initial network, then adjust the thresholds accordingly after assessing the network quality. Suggestions for alterations in thresholds based on the resulting network assessment properties can be found in **Table 2**.

### 9. Assessing Network Quality

**Usage**

~~~
  python ./analysis/assess_network.py --file <network file>
~~~
**Example command**

~~~
  python ./ analysis/assess_network.py --file ./project_folder/out-
  put/network_output/correlations_bw_signif_measurables.csv
~~~
**Inputs**

`--file: correlations_bw_signif_measurables.csv` file created with to_csv.py
**Outputs**

- `network_quality_assessment.txt`: Contains the calculations (also sent to standard output) on the quality of the reconstructed network. Outputs to the same directory the input file is stored in.
- `network.pickle`: A pickled file containing the network. Used as input to future steps. Outputs to the same directory the input file is stored in.

## Network analysis

Once the user has reconstructed a network from the input data, they can move on to the network analysis stage. In this stage, the user can optionally find clusters (also called subnetworks or modules) of nodes and use alternative tools to determine if these clusters are enriched for particular biological processes. Users can then further interrogate the network to find key regulatory nodes in the network.

### 10 (OPTIONAL)

#### a. Identifying clusters of nodes

One commonly performed technique in network analysis is to identify regions in the network where nodes are clustered together. TkNA allows for the use of two popular algorithms, Infomap^16^ and the Louvain^15^ method, to detect clusters in the network. Examples of node clusters can be found in **Figure 6b**.

Infomap

**Usage**

~~~
  python ./analysis/infomap_assignment.py --pickle <file.pickle>
~~~
**Example command**

~~~
  python ./analysis/infomap_assignment.py --pickle ./pro-
  ject_folder/output/network_output/network.pickle
~~~
**Inputs**

`--pickle: network.pickle` file output by `assess_network.py`
**Output**

- `network_infomap_partition.csv`: CSV file containing the name of the node in column 1 and the subnetwork number it was assigned in column 2.

Louvain

**Usage**

~~~
  python ./analysis/Louvain_partition.py --pickle <file.pickle>
~~~
**Example command**

~~~
  Python ./analysis/Louvain_partition.py --pickle ./pro-
  ject_folder/output/network_output/network.pickle
~~~
**Inputs**

`--pickle: network.pickle` file output by assess_network.py
**Output**

- `network_infomap_partition.csv`: CSV file containing the name of the node in column 1 and the subnetwork it was assigned in column 2.

#### b. Perform functional enrichment analysis for groups of nodes

Following clustering, a user can then perform functional enrichment of either 1) the clusters identified via Infomap/Louvain (or externally via MCODE, CLUSTERVIZ, etc.) or 2) any other predefined groups (e.g., microbes, genes), allowing for a biological interpretation of the nodes in that cluster. Single cell RNA-Seq data can also be used to infer cell type information for the genes in the network. Here, cell type is inferred for a gene based on its presence either as a cell cluster’s conserved gene marker or cell cluster specific differentially expressed gene. Additionally, highest average expression of a gene in a specific cell cluster among all clusters can also be used as a genes cell type identity^33^. These enrichment methods are not part of TkNA codes and will need to be performed using external software. Examples of common functional enrichment methods for different omics data are listed in **Table 3**.

### 11. Finding regulatory nodes in the network

This step calculates topological properties for the networks, subnetworks, and nodes^34^. Network properties that are calculated by TkNA include the number of nodes in the network, number of edges, mean degree, average clustering coefficient, the size (number of nodes) of the giant component compared to the size of the whole graph, the number of connected components, mean closeness centrality, degree assortativity^35^, maximum modularity^15^, network fragmentation^36^, and Freeman centralization^37^. Subnetwork properties include the mean degree of the subnetworks/clusters (either pre-defined by the user or identified in the optional step 10) in the network. Finally, calculated node properties include degree, clustering coefficient, closeness centrality, eigenvalue centrality, betweenness centrality, number of second neighbors, and bipartite betweenness centrality (BiBC)^17^. Two of those properties, degree and BiBC, are used to predict nodes with potential regulatory roles in the network. Degree is the number of other nodes a single node connects to. Accordingly, high degree nodes called hubs control the biological pathway/cluster to which they belong.

BiBC is a measure of the bottleneck-ness of a node (i.e., which node serves as the best ‘bridge’) between two pre-defined groups^17^; it is essentially betweenness centrality restricted to particular subsets of the network (i.e., two subgraphs/clusters of biological interest). A node with a high BiBC likely mediates the communication between the selected pathways/clusters. To compute BiBC, the user will need to provide a list of nodes, as well as which cluster they belong to in the network. If the user wishes to perform the BiBC calculation between types of nodes (for example, genes and microbes), they can supply the original sample mapping file. Alternatively, if the user wishes to use the clusters identified with Infomap or Louvain (Step 10, optional), they can supply the output of Step 10 and select two group numbers from the second column as their clusters to use for the BiBC calculation. Examples of nodes with a high degree and a high BiBC can be seen in **Figure 6a**.

**Usage**

~~~
  python ./analysis/calc_network_properties.py --pickle
  <file.pickle> --bibc --bibc-groups <choice> --bibc-calc - type
  <choice> --node-map <file.csv> --node-groups <group 1> <group 2>
~~~
**Example command**

~~~
  python ./analysis/calc-network-properties.py --pickle ./pro-
  ject_folder/output/network_output/network.pickle --bibc --bibc-
  groups node_types --bibc-calc-type rbc --node-map ./pro-
  ject_folder/input/map_file.csv --node-groups micro pheno
~~~
**Inputs and arguments**

`--pickle: network.pickle` file created with `assess_network.py`
`--bibc`: Flag for whether to compute Bipartite Betweenness Centrality (BiBC). This is highly recommended and also required for future steps
`--bibc-groups`: Choice for what to compute BiBC on, either distinct groups (node_types) or on the two most modular regions of the network (found using the Louvain method)
`--bibc-calc-type`: Choice for whether to normalize based on the number of nodes in each group (rbc) or not (bibc)
`--node-map`: CSV file containing the name of nodes in the first column and the type of the node (gene, phenotype, microbe, etc.) in the second column
`--node-groups`: Required if `node_types` is specified for `--bibc-groups`. It’s the two groups of nodes to calculate BiBC/RBC on. The types must be present in the `--node-map file`
**Outputs**

- `network_properties.txt`: Tab-delimited .txt file of calculated network properties
- `subnetwork_properties.txt`: Tab-delimited .txt file of calculated subnetwork properties
- `node_properties.txt`: Tab-delimited .txt file of calculated node properties

## Shortest path calculations

The user can optionally then calculate the distance path between two pathways (subnetworks/clusters) by calculating the shortest path between each member of those clusters and taking the average of those values. Pathways closer to one another potentially interact more than those that are further away.

**Usage**

~~~
  python ./analysis/find_all_shortest_paths_bw_subnets.py --network
  <file.pickle> --node-map <map.csv> --node-groups <group1>
  <group2>
~~~
**Example command**

~~~
  python ./analysis/find_all_shortest_paths_bw_subnets.py --network
  ./project_folder/output/network_output/network.pickle --node-map
  ./project_folder/input/map_file.csv --node-groups gene pheno
~~~
**Inputs**

`--network: network.pickle` file output by `assess_network.py`
`--node-map`: Mapping file (CSV) of nodes and their subnetworks
`--node-groups`: The two groups in the mapping file you want to find the shortest paths between
**Output**

- `shortest_path_bw_<group1>_and_<group2>_results.csv`: CSV file containing the name of each node in each pair in columns 1 and 2, as well as the shortest path length between that pair in column 3 and the number of shortest paths for the pair in column 4

By combining the information about shortest path and BiBC, users can infer interactions between pathways (subnetworks/clusters) as well as causal relations between them, as has been done in our recent study^12^.

## Estimating probability to find top nodes (degree, BiBC) randomly

The next step in TkNA is to determine the likelihood of finding the top nodes (degree, BiBC) randomly in the reconstructed network by comparing to size-matched random networks with the same number of nodes and edges^10^. Later on in the pipeline, the user can visualize how top nodes of their reconstructed network compares to the top nodes of these random networks through the creation of a 2D density plot (**Figure 5d**).

### 12. Creating random networks

This step creates all the random networks and saves them to be analyzed in the next step. Note that due to the complexity of the BiBC calculation, if your network is very large (thousands of nodes and tens of thousands of edges) this step (as well as the next step) can take up to several days. Therefore, we recommend not running this step interactively and instead submitting to a server. If that is not possible, the creation and analysis of random networks can be skipped as it is not required for the visualization of the BiBC and degree calculations. However, **Step 17 “Create 2D density plot”** cannot be run if the random network steps are skipped.

**Usage**

~~~
  python ./random_networks/create_random_networks.py --template-
  network <file.pickle> --networks-file <file.zip>
~~~
**Example command**

~~~
  python ./random_networks/create_random_networks.py --template-
  network ./project_folder/output/network_output/network.pickle --
  networks-file ./project_folder/output/network_output/all_ran-
  dom_nws.zip
~~~
**Inputs**

`--template-network`: The pickled network file output by `assess_network.py`
`--networks-file`: .zip folder to output all random networks to
`--num-networks`: optional; number of random networks to create
**Outputs**

- A single .zip file containing all created networks

### 13. Analyze random networks

Each randomly generated network is then analyzed to calculate the degree and BiBC of each node in the network.

**Usage**

~~~
  python ./random_networks/compute_network_stats.py --networks-file
  <file.zip> --bibc-groups <choice> --bibc-calc-type <choice> --
  stats-file <file.zip> --node-map <file.csv> --node-groups
  <group1> <group2>
~~~
**Example command**

~~~
  python ./random_networks/compute_network_stats.py --networks-file
  ./project_folder/output/network_output/all_random_nws.zip --bibc-
  groups node_types --bibc-calc-type rbc --stats-file ./pro-
  ject_folder/output/network_output/random_network_analysis.zip --
  node-map ./project_folder/input/map_file.csv --node-groups gene
  pheno
~~~
**Inputs**

`--networks-file`: .zip file created with `create_random_networks.py` that contains all random networks previously created
`--bibc-groups`: Group nodes for BiBC based on type or modularity --bibc-calc-type: Compute raw BiBC or normalize (rbc)
`--stats-file`: .zip file to output the network stats to
`--node-map`: CSV file mapping nodes to their types. Required if `node_types` is specified for `--bibc-groups`.
`--node-groups`: Two types of nodes to use for BiBC grouping. Required if `node_types` is specified for `--bibc-groups`.
**Outputs**

- A single .zip file with degree/BiBC results of all random networks

### 14. Condense random network results into a single output file

The results of the analysis are then summarized in a single output file containing the node with the highest degree value and its associated BiBC (or vice versa) in the each of the networks.

**Usage**

~~~
  python ./random_networks/synthesize_network_stats.py --network-
  stats-file <file.zip’> --synthesized-stats-file <file.csv>
~~~
**Example command**

~~~
  python ./random_networks/synthesize_network_stats.py --network-
  stats-file ./project_folder/output/network_output/random_net-
  work_analysis.zip --synthesized-stats-file ./project_folder/out-
  put/network_output/random_networks_synthesized.csv
~~~
**Inputs**

`--network-stats-file`: .zip file created with `compute_network_stats.py`
`--synthesized-stats-file`: Name of the CSV file that will be created
**Outputs**

- A single .csv file that contains the top node, sorted first by BiBC and then by Node_degrees (unless otherwise specified with `--flip-priority`), for each of the random networks

## Plot results

### 15. Create degree distribution and dot plots

In this step, the user specifies which two node properties they would like to visualize for the nodes of the network, as well as how many top nodes to focus on and label. Additionally, the `--top-num-per-type` argument is used to specify how many top nodes per data type to zoom in on and label in the resulting plots. Examples of these visualizations can be seen in **Figure 7**.

**Usage**

~~~
  python ./visualization/dot_plots.py --pickle <file.pickle> --
  node-props <file.txt> --network-file <file.csv> --propx BiBC --
  propy Node_degrees --top-num <integer> --top-num-per-type <integer>
~~~
**Example command**

~~~
  python ./visualization/dot_plots.py --pickle ./pro-
  ject_folder/output/network_output/network.pickle --node-props
  ./project_folder/output/network_output/node-properties.txt --net-
  work-file ./project_folder/output/network_output/network_out-
  put_comp.csv --propx BiBC --propy Node_degrees --top-num 5 --top-
  num-per-type 3
~~~
**Inputs**

`--pickle`: pickled file created with `assess_network.py`
`--node-props: node_properties.txt` file created with `calc_network_properties.py`
`--network-file: network_output_comp.csv` file created with `to_csv.py`
`--propx`: Node property to plot on X-axis
`--propy`: Node property to plot on Y-axis.
`--top-num`: Number of nodes you want to zoom in to on the property v property plot
`--top-num-per-type`: The number of nodes to plot for each data type when zoomed in on the plot
**Default outputs**

- `degree_distribution_dotplot.png`: Distribution of the number of nodes which each degree in the network
- `<propx>_v_<propy>_distribution.png`: A dot plot of user-specified node properties
- `<propx>_v_<propy>_distribution_<node_type>_nodes_only.png`: Same as previous plot, but with just the nodes from each data type. There will be one plot produced for each data type
- `<propx>_v_<propy>_distribution_top_<top-num>_nodes.png`: Same as the second plot, but zoomed in on the top nodes
- `<propx>_v_<propy>_distribution_top_<top-num-per-type>_nodes_<data_type>_only.png`: same as third plot, but zoomed in on the top nodes per data type.

### 16. Create abundance plots

This step creates abundance/expression plots of the top nodes in the network, which are the same nodes that were labeled in the previous step. These figures can be grouped according to class (called ‘Treatment’ in the program) or Experiment.

**Usage**

~~~
python ./visualization/plot_abundance.py --pickle <file.pickle> -
-abund-data <list of files> --metadata <list of files> --color-
group <choice> --x-axis <choice>
~~~
**Example command**

~~~
  python ./visualization/plot_abundance.py --pickle ./pro-
  ject_folder/output/network_output/inputs_for_down-
  stream_plots.pickle --abund-data ./project_folder/input/Expt1.csv
  ./project_folder/input/Expt2.csv --metadata ./project_folder/in-
  put/Expt1_meta.csv ./project_folder/input/Expt2_meta.csv --color-
  group Treatment --x-axis Experiment
~~~
**Inputs**

`--pickle: inputs_for_downstream_plots.pickle` file output by `dot_plots.py`
`--abund_data`: List of data files containing expressions/abundances
`--metadata`: List of metadata files containing Experiment/Treatment columns
`--color-group`: Variable to color the plot by
`--x-axis`: Variable you wish to group the data by on the x-axis
**Outputs**

- One boxplot for each of the top nodes (found in dot_plots.py) as well as additional plots if specified with the optional argument `--nodes-to-plot`

### 17. Create 2D density plot

The last step of TkNA is to create a 2D density plot using the random network results from step 14. This step will take the nodes that were labeled in step 15 and plot them on top of the density plot. It also labels the nodes with the names and probability of a randomly generated network having a node with a higher degree and BiBC than each plotted node.

**Usage**

~~~
python ./visualization/plot_density.py --rand-net <file.csv> --
pickle <file.pickle>
~~~
**Example command**

~~~
  python ./visualization/plot_density.py --rand-net ./pro-
  ject_folder/output/network_output/random_networks_synthesized.csv
  --pickle ./project_folder/output/network_output/inputs_for_down-
  stream_plots.pickle
~~~
**Inputs**

`--rand-net`: file output by `synthesize_network_stats.py`
`--pickle: inputs_for_downstream_plots.pickle` file output by `dot_plots.py`
**Default outputs**

- `density_plot_with_top_nodes_from_dotplots.png`: contour plot with the top nodes (found in `dot_plots.py`) from the real reconstructed network overlaid on top
- `density_plot_with_top_<data_type>_nodes_only.png`: Same as previous, but contains just one data type per output file

## Applications of the method

Although this methodology was originally designed to uncover the interactions between different taxonomic kingdoms, this approach is also used for the analysis of more general types of multiomics data. For example, TkNA can be used to analyze the interaction between genetic and transcriptional data, metabolites, proteins, and phenotypes, as well as various omics data from different organs from the same host.

## Comparison with other methods

The TkNA approach relies on performing meta-analysis across several experiments to identify robust patterns of fold-change and correlations across multiple cohorts. By default, it uses Fisher’s method to combine p-values from several independent tests. Other general R packages (e.g., *meta^38^, netmeta^39^, mixmeta^40^)* offer several methods for meta-analysis that can be used along with the principles of TkNA. Additional R packages (e.g., *MixOmics^41^, MOFA2^42,43^*, iClusterPlus^44^) use sophisticated statistical methods to combine multiple -omics data measured from the same patients. However, their application to several cohorts simultaneously or to unique -omics data (where the data composition or distributional assumptions are not met) can be challenging. TkNA provides a framework to achieve simultaneous integration of multiple -omics types and cohorts.

While many tools claim a gene or microbe to be “important” based on association analysis, TkNA specifically identifies <u>causal</u> relationships rather than only associations.

One of the most popular approaches to determine causality thus far is an application of the Mendelian Randomization^45,46^ (MR) method for the inference of microbes causally contributing to a given host trait. Although this method has a robust theoretical framework, it has two main drawbacks. First, because it is based on differences in causal signals between alleles present in the normal human population, it requires huge sample sizes (e.g., thousands, akin to GWAS) that are often not available, particularly when working with mice. Second, allelic differences in humans explain a very small proportion of microbiome variability, which then limits the approach’s utility to a small handful of microbes clearly regulated by common polymorphisms.

More recently, a variety of regression approaches to causal inference in microbiome experiments have been developed, most of which are based on (linear) structural equation models, which have been around in one form or another for more than a century. The Sparse Microbial Causal Mediation Model (SparseMCMM)^47^ is a recent example. While promising, these group of methods have not been tested in new studies that would experimentally validate model inferences. These approaches deal with the large number of possible covariates (host genes or genetic markers) via either regularization or PCA-type transformation to a reduced set of covariates, potentially making interpretation of gene mediators difficult. Finally, these methods ignore any structure to the covariates (gene-gene interactions) or the microbes (taxa-taxa interactions).

## Experimental design

TkNA requires collection of data from at least two classes (e.g., disease, control). We recommend using at least five biological replicates per sample group when several independent datasets are available. However, ten replicates would be optimal. In regard to statistical consideration, we strongly recommended using two or more independent cohorts/experiments/datasets, which allow for the implementation of meta-analysis ensuring the robustness of inferences from TkNA. We recommend using a standard meta-analysis approach that combines data of the same type from different experiments to identify differentially abundant features (nodes in the network) and correlations from the same class (e.g., “disease”) for establishing edges in the network. However, if the number of available datasets is limited, TkNA allows one to determine edges common across different classes (e.g., “disease”, “control”), either from the same or separate experiments, (see Step 7 in the procedure) (**Figure 8**). This approach increases the power of network reconstruction. However, it is efficient under the strong assumption that differences between classes (e.g., “disease” and “control”) are predominantly limited to the levels of measured variables (e.g., genes, metabolites, etc.) while there are very few edges that are reliably different between networks reconstructed from different classes.

Based on our experience, this assumption holds true for host data such as gene expression and metabolomics, but usually is <u>unreasonable</u> for microbiome data. However, even for host genegene interactions this approach still comes with the cost of losing a few edges specific for individual classes (e.g., relationships that are only present in the disease class). Accordingly, some information related to the biological mechanism underlying a transition from one class to another (e.g., from healthy to disease) might be lost^48–50^.

Inferred pathways and top nodes are biologically informative under any circumstances. However, for causal inference, if additional information allows for the implementation of an “instrumental variable”^51^, the directionality of interactions between kingdoms or pathways can be also established. For example, when studying the effects of antibiotics on a host, it is a reasonable assumption that these effects are mostly mediated by microbes^8,13^; thus microbes in the network can serve as instrumental variables in this case.

## Expertise needed to implement the protocol

The target audience for this protocol is researchers working in the host and/or microbiome field with limited computational and statistical expertise. Our method can be used in biological and biomedical research across **<u>multiple disciplines</u>** ranging from the establishment of new cellular and molecular targets of treatment to fundamental biological questions. This protocol does not require programming expertise; however, users should be comfortable running programs from the command line in the Unix environment and should learn a bit about the JSON file format in order to be able to customize and modify the program’s options.

## Limitations

A basic understanding of statistics is expected from the user to understand the suitability of the software to their experimental setup and any inherent limitations of the approach. For example, Fisher’s method is the default method implemented by this software for combining p-values from independent tests with the same null hypothesis. However, if the same control group is used in calculating the effects of different treatments (or over time), the effect sizes are non-independent. Since the tool cannot automatically detect this dependence and will proceed with the analysis, the user will need to realize that alternative methods to conduct a meta-analysis with non-independent effect sizes^52^ will be more suitable in this case.

The only aspect of the TkNA approach that can be considered as a critical limitation is related to the fact that it requires an appropriate study design for better inference of causal structure of the investigated biological process (this is not unique to TkNA but would be a limitation of any causal inference approach). For example, to define the directionality of a process, as with any other causal inference application (including those mentioned above), it requires study designs that would confer with the assumptions needed for mediation analysis^53^ or for instrumental variable analysis^51^. Since most standard biomedical experiments are designed as “double blind randomized studies” or contain an instrumental variable, TkNA is very powerful for causal inference from multi-omic data.

## Materials

### Required hardware

Memory usage and storage space are primarily correlated with the number of correlations that pass the filters (i.e., the number of edges in the reconstructed network). They are therefore also indirectly correlated with the number of variables that pass the filters for differential expression. As a baseline, 8GB of memory and 10GB of free storage is recommended, but more may be required by larger networks (**Figure 9**).

**Figure 9.**
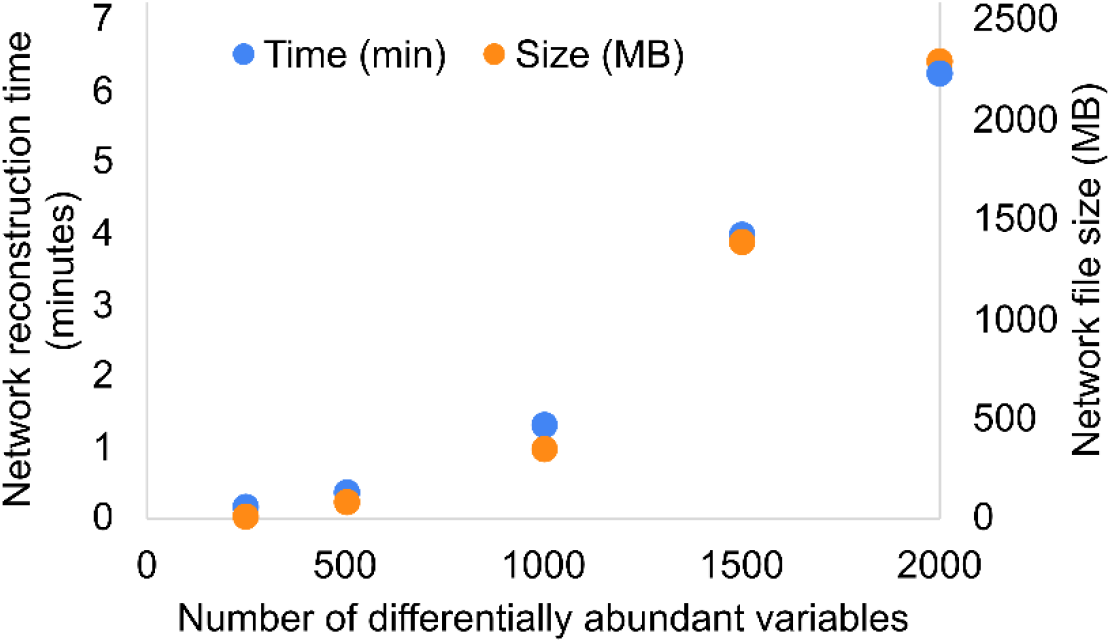
Amount of time (primary axis) and storage space (secondary axis) required to reconstruct a network as a function of the number of differentially abundant variables.

### Benchmarking

Considering the exponential increase in the number of correlations required as the number of differentially abundant variables linearly increases, we ran the network reconstruction section of the TkNA pipeline using various numbers of differentially abundant variables (between 250 to 2000) (**Figure 9**). The reconstruction was performed using 8 cores and 16 GB of RAM. The approximately 2,000,000 correlations required for the largest of these datasets took less than 7 minutes to run.

### Required software and modules

- Miniconda with python 3.8.10 -https://docs.conda.io/en/latest/miniconda.html
- git (as well as a GitHub account) -https://github.com/git-guides/install-git
- Cytoscape - https://cytoscape.org/download.html
- All required modules are in the requirements.txt file in the repository. See step 3 in “Software setup” for how to create a virtual environment and install all the necessary modules to run TkNA.
- Text editing software such as Notepad++, Atom, BBEdit, Gedit, and/or a source-code editor (recommended) such as Visual Studio Code (also known as VS Code) for creating the input files. VS Code aids in the process of writing JSON files.

### Troubleshooting

While this pipeline was tested extensively to ensure compatibility across different environments, it’s not possible to account for all the possible errors one can run into. We have compiled a list below of the most common errors one may run into when running the TkNA pipeline.

- Error: No module named____

- Solution: make sure all the modules and the correct versions are installed from the requirements.txt file.
- Error: Figure legend does not have colors labeled, but instead have the data points labeled in the legend

- Solution: make sure all the modules and the correct versions are installed from the requirements.txt file. Specifically, check the matplotlib and seaborn versions.

### Anticipated results

Through using TkNA, a user should expect to be able to reconstruct and interrogate a network from multi-omic (or single-omic) data. Users can then identify key players (microbes, genes, etc.) that are responsible for the communication between subnetworks. These subnetworks can either be pre-identified by the user or found using community detection algorithms via the Louvain or Infomap algorithms. Functional enrichment of these subnetworks can be performed using external software (not part of TkNA). The nodes that have a high degree and/or high BiBC are key regulatory nodes that contribute to the interaction between these enriched pathways.

Additionally, these findings can be visualized through the use of the TkNA software, including plots for the top nodes based on BiBC and degree, as well as abundance plots of these top nodes and a 2D density plot comparing these top nodes to the top nodes of randomly generated networks. Examples of this data (including all necessary input files and expected output files) for easy-to-understand toy network can be found on the GitHub repo at https://github.com/CAnBioNet/TkNA/tree/main/example_datasets_and_commands/toy_network. We have also uploaded more complex data that includes both microbiome and phenotypic data to https://github.com/CAnBioNet/TkNA/tree/main/example_datasets_and_commands/microbiome_and_phenotype. All input and expected output files are included for that dataset as well.

## Code availability

The TkNA pipeline is publicly available at https://github.com/CAnBioNet/TkNA.

## Contributions

NS, AM conceived the original version of transkingdom network analysis.

NKN, MM, RRR, AKD, NS, GT, KB, AM designed current TkNA framework.

NKN, MM implemented the coding part of TkNA workflow.

RRR, JP prepared parts of TkNA workflow which requires additional software.

AMB, JWP, SSP performed the validation.

NKN, RRR, AMB prepared the simulated data.

RRR prepared the experimental data. NKN, AMB prepared the figures.

NKN, MM, RRR, JP wrote the paper.

AMB, JWP, SSP, JP, AKD, NS, GT, KB, AM edited the paper.

NS, GT, KB, AM supervised various aspects of this study.

## Acknowledgements

The funding that supports this work is AI157369 (AM), DK103761 (NS), DK107603 (AM) and BC011153 (GT). SP and MM were supported by the summer fellowships from the College of Pharmacy at Oregon State University.

## Notes

### Competing Interest Statement

The authors have declared no competing interest.

### Summary of Updates

An additional figure was added (Figure 4) to demonstrate how a network is reconstructed; Figure 6 was added to detail examples of network analysis techniques; An additional plot was added to the previous Figure 5 (now Figure 7) to show an example degree distribution plot; Figure 8 was added to explain possible ways of performing meta-analysis using multiple experimental classes; Figure 9 was added to show the time and storage required for reconstructing networks of various sizes; Text modified throughout the manuscript for clarity; Troubleshooting and Anticipated results sections also added

